# Efficient 5-OP-RU-induced enrichment of Mucosal-associated invariant T cells in the murine lung does not enhance control of aerosol *Mycobacterium tuberculosis* infection

**DOI:** 10.1101/2020.08.19.258509

**Authors:** Charles Kyriakos Vorkas, Olivier Levy, Miroslav Skular, Kelin Li, Jeffrey Aubé, Michael S. Glickman

**Author notes:** Correspondence to Michael Glickman MD, 1275 York Ave, New York, NY 10065.

## Abstract

Mucosal-associated invariant T (MAIT) cells are an innate-like T cell subset in mammals that recognize microbial vitamin B metabolites presented by the evolutionarily conserved MHC I-related molecule MR1. Emerging data suggest that MAIT cells may be an attractive target for vaccine-induced protection against bacterial infections because of their rapid cytotoxic responses at mucosal services to a widely conserved bacterial ligand. In this study, we tested whether a MAIT cell priming strategy could protect against aerosol *Mycobacterium tuberculosis* (*Mtb*) infection in mice. Intranasal co-stimulation with the lipopeptide TLR 2/6 agonist, Pam2Cys (P2C), and the synthetic MR1 ligand, 5-OP-RU, resulted in robust expansion of MAIT cells in lung. Although MAIT cell priming significantly enhanced MAIT cell activation and expansion early after *Mtb* challenge, these MAIT cells did not restrict *Mtb* bacterial load. MAIT cells were depleted later in infection, with decreased detection of granzyme B^+^ and IFNγ^+^ MAIT cells relative to uninfected P2C/5-OP-RU-treated mice. Decreasing the infectious inoculum, varying the time between priming and aerosol infection, and testing MAIT cell priming in NOS2 deficient mice all failed to reveal an effect of P2C/5-OP-RU induced MAIT cells on *Mtb* control. We conclude that intranasal MAIT cell priming in mice induces early MAIT cell activation and expansion after *Mtb* exposure, without attenuating *M. tuberculosis* growth, suggesting that *Mtb* evades MAIT cell-dependent immunity.

## Introduction

Mucosal-associated invariant T (MAIT) cells are an evolutionarily conserved innate-like T cell subset in mammals that recognize microbially-derived Vitamin B intermediates presented by the monomorphic MHC I-related molecule, MR1 (1, 2). Unlike conventional T cells, MAIT cells are activated within hours of antigen recognition (3, 4) and licensed to kill bacterially infected cells (5, 6). MAIT cells also secrete cytokines that recruit accessory immune cells during bacterial infection (7–9). These characteristics suggest that enhancing MAIT cell numbers or function may be an attractive pan-bacterial vaccine approach (10).

MAIT cell responses have been documented in a number of infectious and non-infectious inflammatory diseases (2, 11), including the leading infectious cause of death globally, tuberculosis (TB)(3, 12–19). Peripheral blood MAIT cells contract during active TB disease (13, 17, 18, 20) and are activated during acute exposure in healthy household contacts (3, 15). Limited studies of lung pathology in TB patients suggest that MAIT cells may be recruited to the lung during active infection (21, 22). Better understanding of the role of MAIT cells in host defenses against *Mtb* in vivo will be critical to assessing their contribution to protective immunity. Yet, few studies have examined the role of MAIT cell responses using in vivo animal models of tuberculosis (23–25). Results from previous work are consistent with limited MAIT cell activation and proliferation after *Mtb* infection relative to significant expansion of *Mtb* antigen-specific T cells.

In vivo modeling of MAIT cell responses against *Mtb* have been limited by low MAIT cell abundance in specific pathogen free (SPF) mice (<1% of T cells) relative to the generally more abundant MAIT cell populations observed in humans (<1-18% of T cells) (2, 26, 27). Despite low abundance of MAIT cell populations in SPF mice, several recent studies demonstrate that intranasal priming with bacteria or synthetic MR1 ligands can induce robust MAIT cell accumulation in the lung (27–31). Murine pulmonary MAIT cells significantly expand after intranasal inoculation of *Salmonella typhimurium*, *Francisella tularensis* or *Legionella longbeacheae* in an MR1-dependent manner (29, 30, 32, 33). Importantly, murine pulmonary MAIT cells also expand after intranasal priming with synthetic MR1 ligand 5-OP-RU + TLR ligand co-stimulation, but not with 5-OP-RU or TLR ligands alone (29). This co-stimulatory priming strategy is protective against murine pulmonary *L. longbeacheae* infection (30). Taken together, these data support the hypothesis that MR1 ligand induced MAIT cell populations may protect against other intracellular pulmonary pathogens, including *Mycobacterium tuberculosis* (*Mtb*) (34, 35). Here, we assessed a MAIT cell priming strategy in the murine model of aerosol *M. tuberculosis* infection.

In the following experiments, we tested the hypothesis that MAIT cell priming with synthetic lipopeptide TLR2/6 agonist, Pam2Cys (P2C) + 5-OP-RU co-stimulation can attenuate aerosol *Mtb* infection in mice. We demonstrate that vaccination with P2C/5-OP-RU prior to *Mtb* challenge enhanced MAIT cell activation and expansion by 7 days after infection. Despite significant recruitment to lung after acute bacterial exposure, MAIT cells were depleted by week 3 post-infection and no attenuation of bacterial load was observed. Decreasing the infectious inoculum, the time between priming and *Mtb* aerosol challenge, and ablating host NOS2 all failed to reveal a salutary effect of MAIT cell priming. Our study demonstrates that MAIT cell priming enhances MAIT cell number and activation in *Mtb* infected lungs, but does not attenuate the course of infection.

## Results

### MAIT cells respond to acute *Mtb* challenge with limited expansion in lung and mediastinal lymph node during the chronic phase of infection

We first infected wild type C57/Bl6 (WT) from Jackson™ laboratories with “low-dose” (LD) *Mtb* inoculum (50-150 cfu/lung) via aerosol route and quantified MAIT cell abundance and activation by flow cytometry 7 and 21 days following infection. The MAIT cell gating strategy is defined in Supplemental Figure 1a and all MR1-5-OP-RU staining was controlled with MR1-6FP tetramers. As previously reported, MAIT cells are rare in SPF mice (<1%), but can be identified using MR1-5-OP-RU tetramers (26) and were most clearly detected in inguinal lymph nodes in uninfected mice (Figure 1a, b). Seven days following *Mtb* infection, there was no significant expansion of MAIT cell subsets and in fact we observed fewer MAIT cells in spleen and inguinal lymph nodes (Figure 1b) in infected mice with a trend towards depletion in lung in the most prevalent double negative (DN) subset (Figure 1c). This was accompanied by relative enrichment in the minor CD4^+^ subset (Figure 1c). There was also a trend towards increased MAIT cell activation marker expression in infected mice, though this did not reach statistical significance (Figure 1d). By 21 days post-infection, MAIT cells expanded in infected lung and mediastinal lymph node (Figure 1e,f). In lung, the DN MAIT cell subset underwent the most significant expansion, but increased MAIT cell numbers were also noted in CD4^+^ and CD8^+^ MAIT cell subsets (Figure 1g). As expected, non-MAIT T cells significantly expanded in lung at the onset of the adaptive immune response, most notable in the CD4^+^ T cell subset (Figure 1g). Pulmonary MAIT cells from mice infected for 21 days upregulated activation marker CD69 and natural killer receptor NK1.1 relative to uninfected mice (Figure 1h). As expected during the adaptive response to *Mtb* infection, pulmonary non-MAIT T cells strongly downregulated the naive L-selectin marker, CD62L. These data are consistent with previous findings (23–25) demonstrating that MAIT cells respond to acute *Mtb* challenge, but undergo limited expansion.

**Figure 1.**
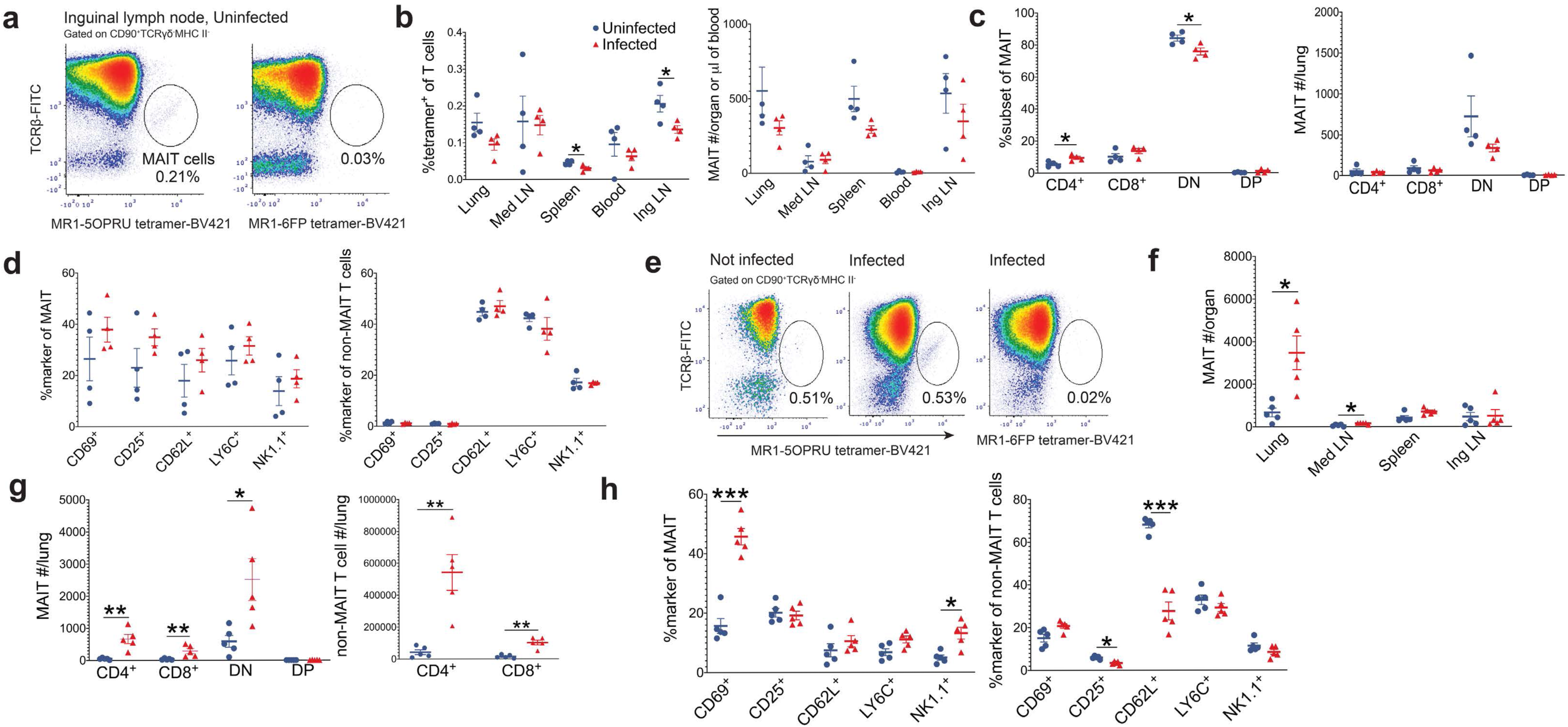
MAIT cells respond to *Mycobacterium tuberculosis* infection in mice. **(a)** Representative flow cytometry density plots from one mouse identifying MAIT cells using MR1-5-OP-RU tetramers in the inguinal lymph node compared to negative control MR1-6FP tetramers. **(b)** Mean % of MAIT cells of CD90^+^ cells +/− SEM and the mean MAIT cell absolute number +/− SEM within tissues of uninfected and *Mycobacterium tuberculosis* (*Mtb*)-infected mice 7 days post-infection. blue=not infected and red=infected. **(c)** MAIT cell CD4^+^ or CD8^+^ co-expression as mean % of total MAIT cells +/− SEM and mean absolute number +/− SEM 7 days post-infection. **(d)** Mean % Activation marker^+^ +/− SEM of pulmonary MAIT cells and non-MAIT T cells 7 days post-infection. **(e)** Representative flow cytometry density plots from one infected mouse 21 days after aerosol challenge and one uninfected mouse identifying pulmonary MAIT cells using MR1-5-OP-RU tetramers. Staining was controlled using MR1-6FP tetramers. **(f)** Mean MAIT cell absolute number +/− SEM in various tissues 21 days post-infection. n=5 mice/group **(g)** Mean absolute number +/− SEM of MAIT cell and non-MAIT cell subsets 21 days post-infection. **(h)** Mean% Activation marker^+^ +/− SEM of MAIT and non-MAIT T cells 21 days post-infection. n= 4-5 mice/group. All data in this figure represent one independent experiment. Statistical analyses were performed using unpaired t-tests with significance level of p<0.05. *p<0.05 **p<0.005 *** p<0.0005

### Intranasal priming with 5-OP-RU and TLR2/6 ligand Pam2Cys co-stimulation induces robust MAIT cell expansion within lung

To ask whether enhancement of MAIT cell numbers or function could affect *Mtb* infection, we adapted a model of MAIT cell priming (29, 30), which is summarized in Figure 2a. For all priming experiments, we used WT C57Bl/6 mice from Taconic™ laboratories due to higher lung MAIT cell numbers detected at baseline relative to mice from other vendors (Supplemental Figure 2a,b). The gating strategy for all subsequent experiments is defined in Supplemental Figure 1b.

**Figure 2.**
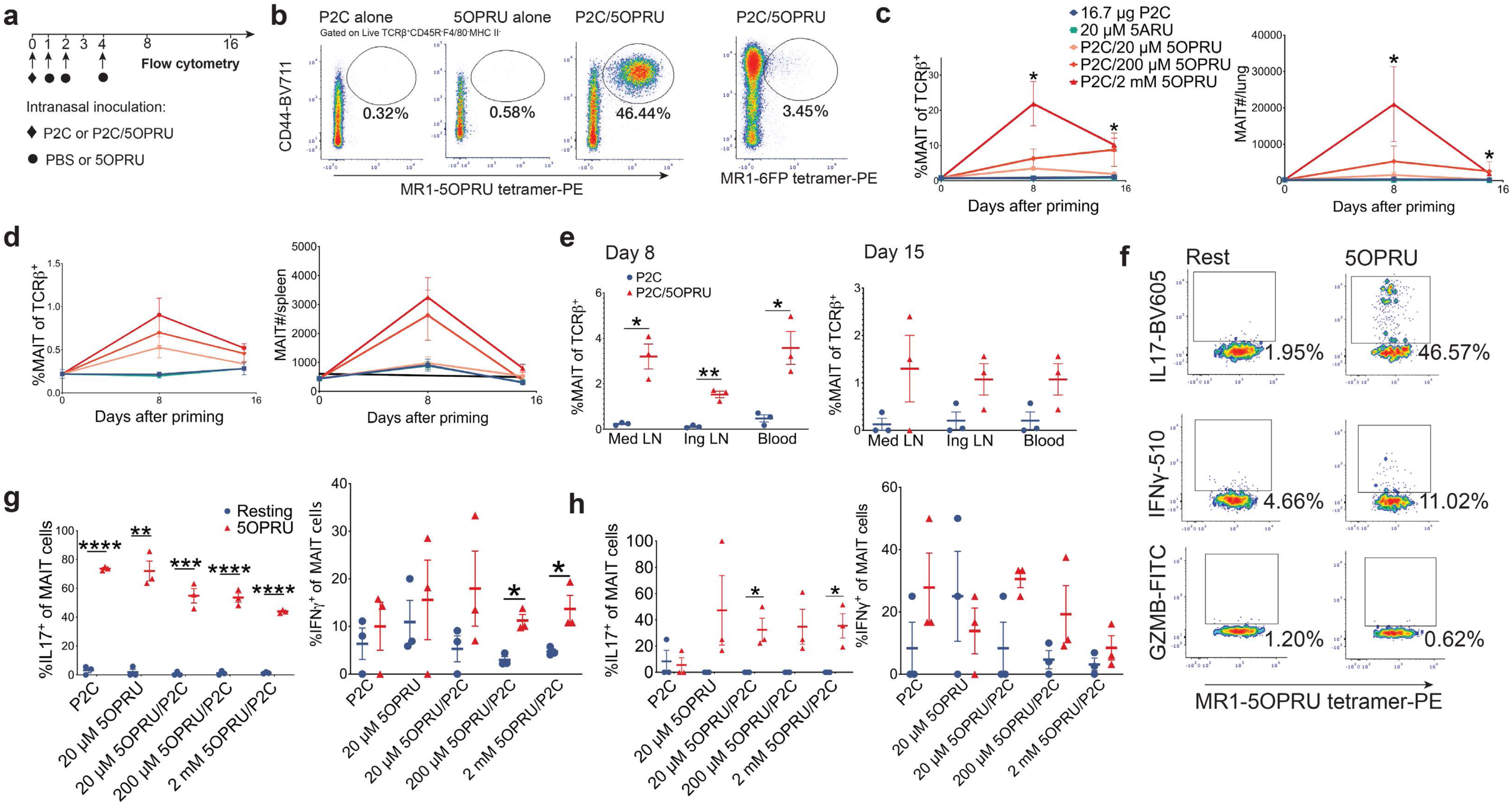
Pulmonary MAIT cells expand after intranasal inoculation with MR1 ligand (5-OP-RU) and Toll-like receptor 2/6 ligand (Pam2Cys) co-stimulation. **(a)** Experimental schematic of intranasal priming model for pulmonary MAIT cell expansion. **(b)** Representative flow cytometry density plots from lungs of three mice 8 days after intranasal priming under different conditions. Pulmonary MAIT cells were identified using MR1-5-OP-RU tetramers. Staining was controlled using MR1-6FP tetramers. **(c,d)** Mean% MAIT cells of T cells +/− SEM and mean MAIT cell absolute numbers +/− SEM measured at baseline then 8 and 15 days after priming under different conditions in lung (c) and spleen (d). Statistically significant results shown compare P2C/2 mM 5-OP-RU with P2C alone. **(e)** Mean %MAIT of T cells +/− SEM in lymph nodes and blood at day 8 or 15 following priming with 2 μM 5ARU/16.7 μg (Pam2Cys) P2C or P2C alone. **(f)** Representative flow cytometry density plots demonstrating intracellular cytokine staining of pulmonary MAIT cells for IL17, IFNγ or granzyme B. MAIT cells were isolated 15 days after intranasal priming with 2 μM 5-OP-RU/16.7 μg P2C and then incubated in vitro for 15 hours at rest or in the presence of 2 μM 5-OP-RU. **(g, h)** Intracellular staining of IL17 and IFNγ in pulmonary (g) and splenic (h) MAIT cells incubated in the same in vitro conditions as described in panel (f). n=3 mice/group. Each panel is representative of three independent experiments. Statistical analyses were performed using unpaired t-tests. *p<0.05 **p<0.005 *** p<0.0005 ****p<0.0001

In vitro, 5-OP-RU alone was sufficient to expand splenic MAIT cells (Supplemental Figure 3a,b). However, in vivo, TLR co-stimulation with 5-OP-RU is necessary for efficient MAIT cell expansion in lung (29) (Fig. 2b-d; Supplementary Figure 3c). First, we confirmed that intranasal administration of MR1 ligand 5-OP-RU + TLR 2/6 ligand Pam2Cys (P2C) induces a dose-dependent MAIT cell expansion in lung (Figure 2b,c) and to a lesser extent, spleen (Figure 2d). In contrast, P2C/5-OP-RU priming via oral gavage did not induce MAIT cell expansion (Supplemental Figure 3c). Transient MAIT cell expansion after intranasal priming was also observed in mediastinal and inguinal lymph nodes as well as blood (Figure 2e), but relatively few MAIT cells were detected in other organs (Supplemental Figure 3d). Functionally, murine pulmonary MAIT cells strongly upregulated IL17 and to lesser degree IFNγ after ex vivo re-stimulation with 5-OP-RU (Figure 2f, g), whereas only IL17 was upregulated in splenic MAIT cells (Figure 2h). These experiments demonstrate efficient induction of pulmonary MAIT cell proliferation and effector functions by exogenous ligand administration and replicate published studies (29, 30).

**Figure 3.**
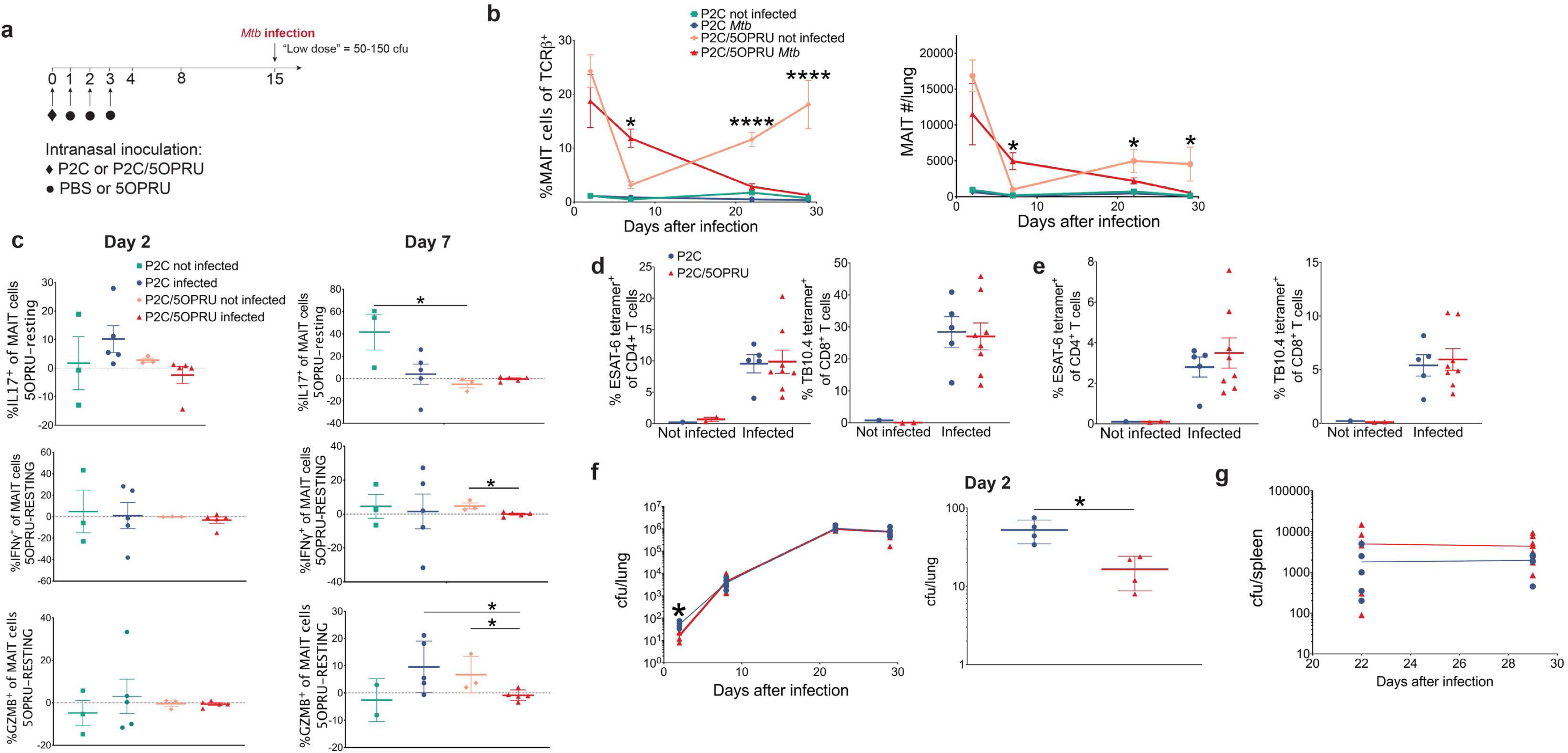
Intranasal priming with P2C/5-OP-RU prior to *Mtb* infection induces early MAIT cell expansion and effector function without impacting bacterial load. **(a)** Experimental schematic of MAIT cell priming prior to *Mtb* infection. **(b)** Mean% MAIT cells of T cells +/− SEM and mean MAIT cell absolute number +/− SEM in lung over time in uninfected and infected mice receiving intranasal inoculation with 16.7 μg of P2C alone or P2C/2 mM 5-OP-RU co-stimulation. Time 0 is the time of aerosol challenge in infected mice. **(c)** MAIT cell cytokine staining in mouse lung from Day 2 and 7 post-infection, measured by the mean % difference between 5-OP-RU and resting conditions +/− SEM after 15 hrs of in vitro re-stimulation. **(d,e)** Mean % ESAT-6- or TB10.4-specific T cells +/− SEM in lung **(d)** or spleen **(e)** at Day 29 post-infection. **(f, g)** *Mtb* colony forming units (cfu) in lung (f) and spleen (g) over time. n=3-8 mice/group. Data in this figure is representative of three independent experiments. Statistical analyses were performed using unpaired t-tests. *p<0.05 ****p<0.0001

### 5-OP-RU + Pam2Cys priming induces early MAIT cell activation and expansion in Mtb-infected lung without attenuating bacterial load

We next tested this MAIT cell priming model as a preventive strategy against *Mtb* infection. Mice were primed via intranasal inoculation and infected with 50-150 cfu/lung via aerosol 14 days after initiation of priming as summarized in Figure 3a. Increased numbers of MAIT cells in lung, both as a percentage of total T cells and in absolute numbers, were detected in P2C/5-OP-RU primed mice at day 7 post-infection compared to either infected P2C controls or uninfected P2C/5-OP-RU mice (Figure 3b), consistent with enhanced MAIT cell activation and expansion during acute infection with vaccination. The inverse trend was observed during the adaptive phase of immunity, where P2C/5-OP-RU treated infected mice had fewer MAIT cells in lung than P2C/5-OP-RU treated but uninfected mice, consistent with depletion of MAIT cells during later phases of infection. By Day 7 post-infection, P2C/5-OP-RU treated mice had blunted MAIT cell IFNγ and granzyme B responses to 5-OP-RU re-stimulation in vitro compared to uninfected mice (Figure 3c).

We next asked whether MAIT cell priming could enhance antigen-specific T cell responses by quantitating ESAT-6 and TB10.4 *Mtb*-specific T cells using MHC tetramers. The gating strategy for antigen-specific T cells is defined in Supplemental Figure 3. No difference in abundance of ESAT-6 or TB10.4 antigen-specific T cells was detected in lung or spleen of P2C/5-OP-RU primed mice by days 22 or 29 post-infection (Figure 3d,e), compared to infected mice treated with P2C alone. Consistent with previous observations (24), antigen-specific T cells greatly outnumbered MAIT cells in lung during chronic infection. Although we observed a small difference in *Mtb* bacterial load immediately after aerosol deposition in P2C/5-OP-RU-treated mice, this difference was not sustained such that bacterial loads in lung or spleen were the same in both groups at later time points (Figure 3f, g).

### Decreasing the *Mtb* inoculum or shortening the time to infection after priming did not enhance MAIT cell mediated control of Mtb

We hypothesized that the “low-dose” *Mtb* inoculum targeting 50-150 cfu/lung was still higher than the physiologic inoculum of natural human infection (36, 37), and could overwhelm early effector functions of MAIT cells. We therefore compared our standard “low-dose” (LD) inoculum of 50-150 cfu/lung with an “ultra low-dose” (ULD) inoculum of 1-10 cfu/mouse (Figure 4a). Using the ULD inoculum, some mice receiving aerosol challenge may remain uninfected. Although we observed fewer cfu in lung 21 days post-infection using ULD versus LD inocula, P2C/5-OP-RU priming again did not decrease bacterial load with either inoculum (Figure 4b).

**Figure 4.**
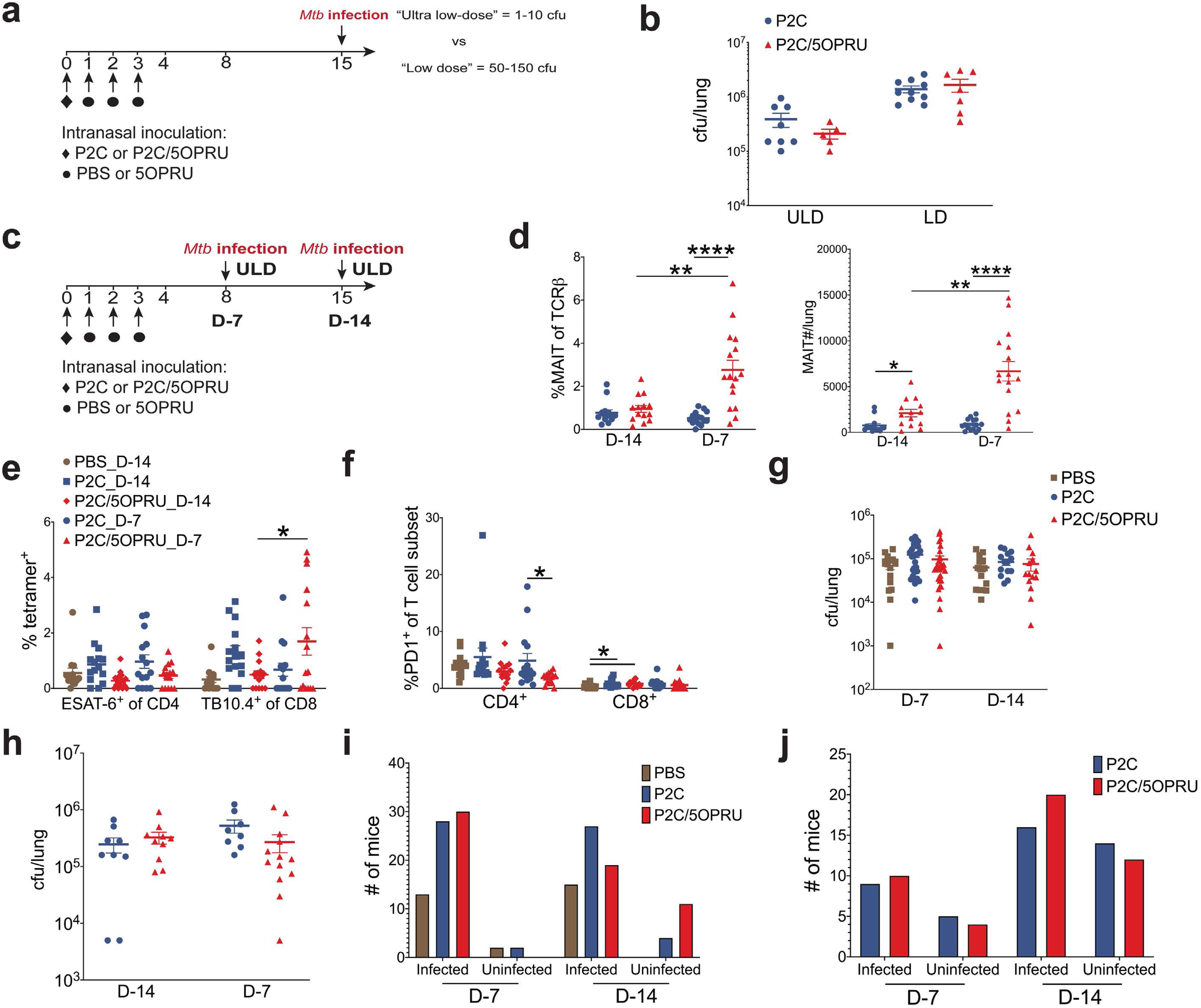
Variation of *Mtb* inoculum or infection schedule did not attenuate Mtb infection in MAIT cell-primed mice. **(a)** Experimental schematic of ultra-low dose (ULD) vs low-dose (LD) *Mtb* infection after MAIT cell priming. **(b)** Mean *Mtb* cfu +/− SEM in lung at 21 days post-infection stratified by inoculum. **(c)** Experimental schematic of two infection schedules. **(d)** Mean % MAIT cells of T cells +/− SEM and mean MAIT cell absolute number +/− SEM in lung 21 days post-infection stratified by infection schedule. **(e)** Mean %ESAT-6 or TB10.4^+^ T cells of CD4^+^ or CD8^+^ T cells +/− SEM 21 days post-infection under different infection schedules. **(f)** Mean %PD1^+^ of CD4^+^ or CD8^+^ T cells +/− SEM 21 day post-infection **(g, h)** Mean *Mtb* cfu +/− SEM at 14 (g) and (h) 24 days post-infection using two infection schedules. **(i, j)** Number of infected and uninfected mice per treatment group after aerosol challenge with ULD inoculum at 14 (i) and (j) 24 days post-infection. Data in panels b and d-f represent one independent experiment. Data from panels g/i and h/j represent 3 and 2 independent experiments, respectively. n=15-30 mice/group. Statistical analyses were performed using unpaired t-tests *p<0.05 *** p<0.0005 ****p<0.0001

Due to the transient kinetics of MAIT cell expansion after priming, we also hypothesized that the timing of infection after P2C/5-OP-RU treatment may impact the efficacy of our vaccination strategy. We subsequently compared two infection schedules using ULD inoculum, conducting aerosol challenge 7 (D-7) or 14 days (D-14) after initiation of priming (Figure 4c). Significantly increased MAIT cell numbers in lung were observed using the D-7 infection schedule (Figure 4d), compared to D-14. This enhanced MAIT cell enrichment was accompanied by a significant increase in TB10.4-specific CD8^+^ T cells and decreased PD1 expression on CD4^+^ T cells (Figure 4e,f) relative to the D-14 infection schedule. However, despite this enhanced MAIT cell and antigen-specific T cell accumulation, P2C/5-OP-RU priming again had no impact on bacterial load (Figure 4g, h). As a subset of mice remained uninfected after ULD *Mtb* aerosol challenge, we compared the fraction of infected mice in each group, reasoning that MAIT cell priming might completely abort infection in some mice. However, we detected no significant differences across treatment groups and timepoints (Figure 4i, j).

We also note that P2C/5-OP-RU priming had no significant effect on the expansion of other immune subsets during infection including non-MAIT CD4^+^ or CD8^+^ T, γδT, B, NK, or myeloid cells (data not shown). *Mtb* cfu did not significantly correlate with absolute number of MAIT cells or any other immune subset in P2C/5-OP-RU treated mice (data not shown).

We also observed that P2C/5-OP-RU priming was associated with decreased intracellular granzyme B and IFNγ in MAIT cells of infected mice on day 14 and 21 post-infection, consistent with functional exhaustion relative to mice treated with P2C alone (Supplemental Figure 5a-c). In the absence of infection, we observed that MAIT cells strongly upregulated the checkpoint inhibitor PD1 with serial P2C/5-OP-RU treatments without any additive effects on proliferation (Supplemental Figure 5d-f). Together, these results demonstrate that priming with synthetic ligand can drive MAIT cells to phenotypic and functional exhaustion.

**Figure 5.**
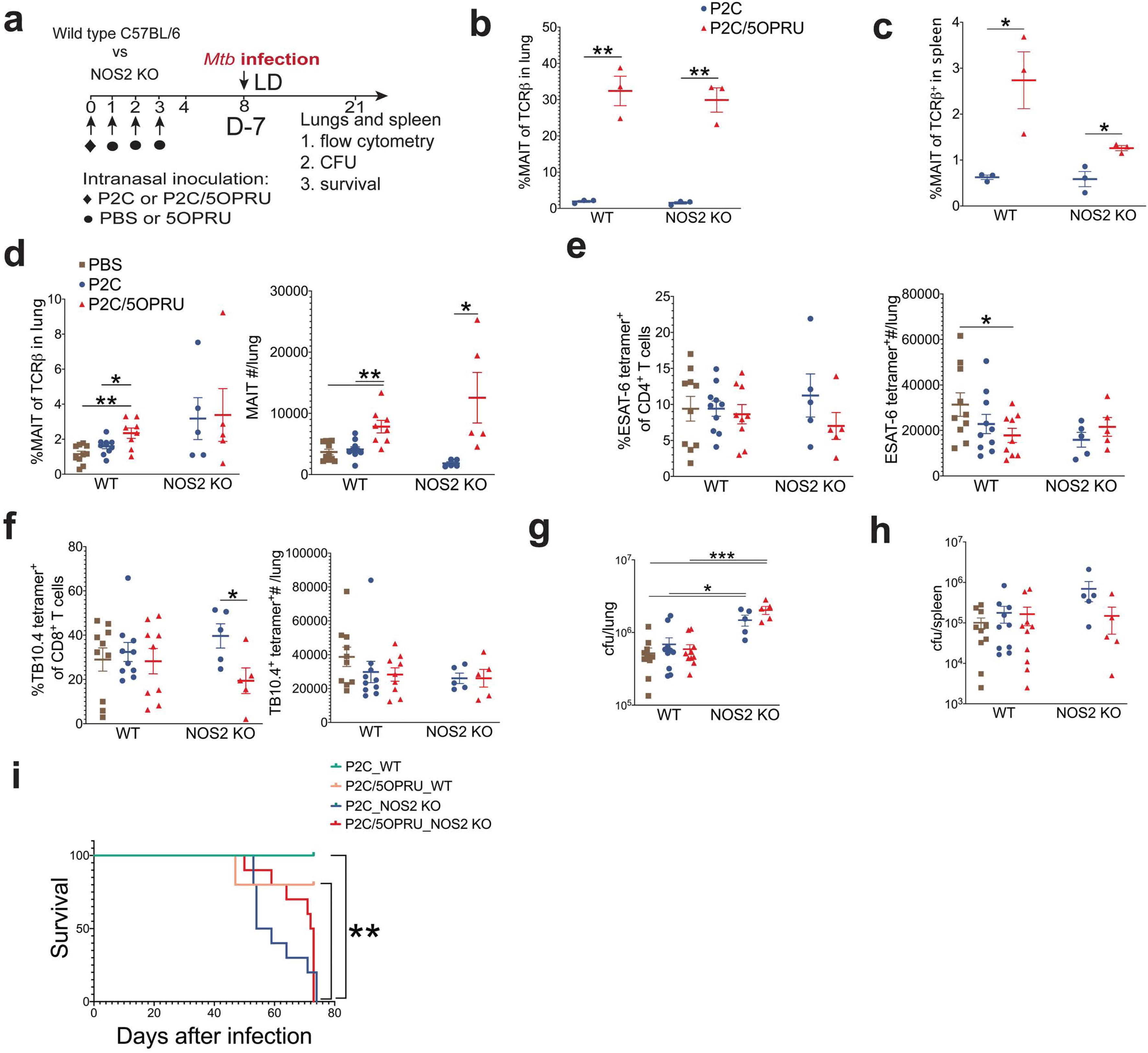
MAIT cell priming does not rescue hypersusceptible iNOS2 deficient mice from lethal infection. **(a)** Experimental schematic of MAIT cell priming prior to *Mtb* infection of wild type (WT) C57Bl/6 or inducible nitric oxide synthase 2 (iNOS2) knock-out (KO) mice. **(b,c)** Mean% MAIT cells of T cells +/− SEM in lung (b) and spleen (c) of uninfected mice 7 days after receiving intranasal priming. n=3 mice/group in panels b and c. **(d)** Mean% MAIT cells of T cells +/− SEM and mean MAIT cell absolute number +/− SEM in lungs 21 days after infection. **(e,f)** Mean% ESAT-6- (e) or TB10.4- (f) antigen-specific T cells of CD4^+^ or CD8^+^ T cells, respectively, +/− SEM 21 days post-infection. **(g, h)** Mean *Mtb* cfu +/− SEM in WT and NOS2 KO mice in lungs and spleen (h) 21 days post-infection. n=5-10 mice/group in panels d-h. **(i)** Survival of *Mtb*-infected WT (n=5 mice/group) and NOS2 KO mice (n=5-10 mice/group). All data in this figure represent one independent experiment. Statistical analyses were performed using unpaired t-tests in all panels except for g where a log rank test was used. *p<0.05 **p<0.005 *** p<0.0005 ****p<0.0001

### MAIT cell priming does not rescue hypersusceptible NOS2 deficient mice from lethal infection

We next hypothesized that MAIT cell priming could attenuate *Mtb* infection in genetically hypersusceptible mice deficient in inducible nitric oxide synthase 2 (NOS2), an enzyme essential to control chronic *Mtb* infection. NOS2 deficient mice lack nitric oxide synthase and succumb to chronic *Mtb* infection within months relative to wild type C57Bl/6 mice that can survive with chronic infection indefinitely despite disseminated disease and high bacterial burden (38, 39). The experimental schematic of infection is presented in Figure 5a. We first confirmed that P2C/5-OP-RU treatment induced equivalent enrichment of MAIT cells in lung in both WT and NOS2 deficient strains in the absence of infection (Figure 5b). MAIT cell enrichment in spleen was decreased in NOS2 deficient mice (Figure 5c). We then infected WT and NOS2 deficient mice with a standard LD *Mtb* inoculum of 50-150 cfu/lung and evaluated immunologic and bacteriologic outcomes 21 days post-infection as well as mortality. Absolute numbers of MAIT cells were enriched in lungs of P2C/5-OP-RU primed mice in both strains (Figure 5d). We observed no significant difference in *Mtb* antigen-specific T cells between treatment groups (Figure 5e,f). As expected, NOS2 deficient mice had approximately 10-fold higher bacterial load than WT mice, but no significant difference was observed in lung bacterial load after P2C/5-OP-RU priming (Figure 5g). Although there was a trend toward enhanced spleen bacterial control in P2C/5-OP-RU treated NOS2 deficient mice, this trend did not reach statistical significance (693,000 v 149,000, p=.18, Figure 5h). All 20 NOS2 deficient mice died or were euthanized as a humane endpoint by ~11 weeks after infection while 9/10 WT mice survived. We observed a longer median survival of P2C/5-OP-RU treated NOS2 deficient mice relative to P2C alone, but this trend also did not reach statistical significance (72.5 v 56.5 days; p=0.18) (Figure 5i).

## Discussion

Emerging evidence from murine infection models of *Francisella* and *Legionella spp* support a protective role for MAIT cells against intracellular pulmonary bacterial pathogens (10, 30, 32, 33). Importantly, MAIT cell priming prior to murine *Legionella longbeacheae* infection significantly attenuated bacterial load and was dependent on MR1, GMCSF and IFNγ, but not on perforin, IL17A or TNFα(30, 32).

In contrast to these studies, prior literature indicates that MAIT cells undergo limited expansion after pulmonary *Mtb* infection in mice and macaques (23–25). This is consistent with our findings: we observed initial MAIT cell depletion from inguinal lymph node, spleen and lung at Day 7 post-infection, likely explained by rapid MAIT cell TCR downregulation and apoptosis following acute exposure (3, 26, 40–42). After 21 days, *Mtb* induced only a mild MAIT cell expansion in lung and mediastinal lymph node when antigen-specific conventional T cells were robustly proliferating (43). Our data, together with previously published observations in humans (18) and animal models (24, 25) suggest that early MAIT cell responses may be suppressed by *Mtb* infection.

Despite this limited expansion, several pieces of evidence suggested that selective MAIT cell priming would enhance antimycobacterial activity. In vitro, MAIT cells can inhibit BCG growth in infected murine macrophages (44). In vivo, transgenic mice over-expressing the murine MAIT cell Vα19 TCR demonstrate decreased *Mtb* lung burden compared to wildtype mice (23). In BCG vaccinated macaques and humans, MAIT cells demonstrate rapid expansion and enhanced IFNγ production(14, 34, 45). In healthy household contacts of TB patients, MAIT cells demonstrate enhanced function in highly exposed donors who remain uninfected (3, 15). Taken together, these data support the hypothesis that direct targeting of MAIT cells prior to *Mtb* challenge may attenuate the course of infection.

We hypothesized that recruiting MAIT cells to lung prior to aerosol *Mtb* challenge could enhance their cytotoxic and helper activity during infection and confer protective immunity. However, our results convincingly demonstrate that intrapulmonary MAIT cell activation and expansion prior to *Mtb* challenge is not sufficient to attenuate bacterial load. Further, we show that while MAIT cells are activated and expand early in P2C/5-OP-RU infected mice, they are subsequently depleted and driven to functional exhaustion. These results mirror observations of peripheral MAIT cell depletion and upregulation of exhaustion markers during active TB infection in humans(12, 13, 17). We also highlight that neither decreasing *Mtb* inoculum nor the time between priming and infection impacted bacteriologic outcomes. Overall, our study findings are consistent with early MAIT cell activation and expansion with priming prior to *Mtb* infection, but no protection against murine aerosol *Mtb* infection.

Our results are consistent with recent work in *Mtb*-infected macaques demonstrating mild enrichment of MAIT cells in bronchoalveolar fluid by 21 days post-infection, a timepoint when *Mtb*-antigen-specific T cells are more robustly expanding in lung (24). In that study, MAIT cells were also sparsely detected in granulomas and did not correlate with *Mtb* bacterial load, suggesting a limited role for MAIT cell immunity against *Mtb* in the non-human primate model.

Importantly, our results strikingly contrast with MAIT cell priming efficacy in a *Legionella* model (30, 32) and prompt further inquiry into why MAIT cells are unable to control *Mtb* infection despite enhanced numbers and effector functions. Specifically, the efficacy of targeted MAIT cell immune therapy may depend upon the capacity of the pathogen itself to induce MAIT cell proliferation as seen with *Francisella* (33) or *Legionella spp* (30, 3*2*), a feature that may reflect the availability of appropriate MR1 presented ligand.

One technical difference between infection models is that sublethal doses of *Legionella longbeacheae* or *Francisella* live vaccine strain (LVS) lead to attenuated infection and eventual clearance by wild type mice, even without P2C/5-OP-RU priming (30, 33). In contrast, we show that even an “ultra low-dose” *Mtb* infection targeting 1-10 cfu/mouse results in chronic infection with high bacterial burdens that cannot be cleared without antibiotics (46, 47). However, we do not believe this alone can explain the distinct MAIT cell vaccination efficacy between models since protection conferred by MAIT cell priming against *Legionella* was observed early after infectious challenge at the peak of bacterial burden.

The microbial mechanisms underlying attenuated MAIT cell responses against *Mtb* infection in animal models remain poorly understood. We speculate that MAIT cells are inefficiently activated by *Mtb*-derived MR1 ligands during in vivo infection. One reason may be that *Mtb* does not provide sufficient quantities of activating MR1 ligands to the host, due to slower replication rates (48, 49), resulting in inefficient MR1 trafficking and presentation in vivo (50). Additionally, *Mtb* could produce inhibitory MR1 ligands that suppress MAIT cell activation. Recent studies suggest that mycobacteria-derived MR1 ligand composition differs from other bacterial species (50, 51). Mass spectrometry of eluates from human tetramers bearing ligands from the evolutionarily-related mycobacterium, *M. smegmatis* detected decreased MR1 activating ligands rRL-6-CH2OH/5-OP-RU and RL-6-M27-OH relative to *E. coli* (51). Moreover, MR1 inhibitory ligands including riboflavin itself and 7,8-didemethyl-8-hydroxy-5-deazariboflavin (FO) were only detected in *M. smegmatis* tetramers, but not *E. coli*(51). Importantly, these differences translated into decreased IFNγ responses of MR1-reactive T cell clones to *M. smegmatis*-derived ligands compared to *E. coli*. Taken together, these data raise the possibility that *Mtb*-derived MR1 ligands may induce a net inhibitory effect on MR1 signaling and suppress MAIT cell activation, despite efficient 5-OP-RU priming prior to infection.

Our results also contrast with the immunotherapeutic efficacy of the other main *Mtb*-reactive innate-like T cell subset: Vγ9;δ2 T cells that respond to *Mtb*-derived phosphoantigens in primates (3, 52, 53). Pulmonary *Mtb* infection alone induces robust Vγ9δ2 T cell proliferation in macaques (54) and intrapulmonary phosphoantigen/IL2 priming prior to *Mtb* challenge enhanced recruitment of *Mtb* antigen-specific T cells to lung and attenuated bacterial load(55). In contrast, P2C/5-OP-RU priming of MAIT cells neither attenuated infection nor significantly enhanced recruitment of *Mtb*-antigen specific T cells.

Emerging data would suggest that adjunctive cytokine stimulation may enhance MAIT cell activity against mycobacterial infection. MAIT cell responses from BCG re-vaccinated human donors were more dependent on IL12/IL18 signaling than MR1(14). In a separate study, enhanced MAIT cell responses from human tuberculous pleural effusions were also found to be cytokine-dependent (21). Importantly, IL23 was shown to be necessary for MAIT cell expansion after murine pulmonary *S. typhimurium* infection(32). In this same study, administration of synthetic 5-OP-RU + IL23 co-stimulation enhanced antimicrobial responses against murine *Legionella* similar to TLR 2/6 ligand+5-OP-RU co-stimulation previously reported (30, 32). MAIT cell protective immunity during *Mtb* infection may require additional activating signals not acquired by P2C + 5-OP-RU priming alone. Future studies should test additional MAIT cell priming approaches against *Mtb*, including cytokine co-stimulation with IL12/18/IL23.

In sum, we demonstrate that intranasal 5-OP-RU and TLR 2/6 co-stimulation prior to aerosol *Mtb* challenge in mice enhances MAIT cell activation and expansion in lung, but is not sufficient to attenuate infection. Our findings stand in contrast with the efficacy of MAIT cell priming observed in a *Legionella* infection model, indicating that the translational potential of MAIT cell priming is likely pathogen-specific. These results will make important contributions to future studies targeting MAIT cells as antimicrobial immunotherapy.

## Acknowledgments

We would like to thank Ashutosh Chaudhry and Alexander Rudensky, Sloan Kettering Institute, MSKCC for valuable advice and discussions while conducting these experiments. We thank Kevin Urdahl, University of Washington for his guidance regarding the “ultra low-dose” inoculum in our *Mtb* aerosol infection model. We also thank Matthew Adamow and Phillip Wong of the MSKCC Immune Monitoring core facility as well as Rui Gardner, Kathy Daniels, Mark Kweens, and Fang Fang of the MSKCC Flow cytometry core facility for their expert consultation. MR1, ESAT-6 and TB10.4 tetramers were produced by the NIH tetramer Core Facility. This work was supported by the Tri-I TBRU, part of the TBRU Network (U19 AI111143), and P30 CA008748. C.K.V. was supported by NIAID K08AI132739.

## Author Contributions

C.K.V. and M.S.G. conceived of the study. C.K.V., O.L, and M.S. conducted the experiments. K.L. and J.A. synthesized 5-A-RU. C.K.V. and M.S.G. analyzed and interpreted the data, and wrote the manuscript with input from all the authors.

## Materials and Methods

### Animals and *M. tuberculosis* infections

Wild-type C57BL/6 were obtained from Jackson laboratories (C57BL/6J #000664; Fig. 1) or Taconic laboratories (B6NTac; Fig. 2-5). Additional wild-type C57BL/6 mice were obtained from Envigo (C57BL/6JRccHsd) and Charles River (C57BL/6NCrl; #027) laboratories (Supplementary Fig. 2). iNOS2 deficient mice (B6.129P2-Nos2^tm1Lau^/J #002609) were obtained from Jackson laboratories (Fig. 5). Female mice were used during *Mtb* infection due to extended duration of co-housing. All mice were housed in an animal biological safety level 3 (ABSL3) vivarium which is fully accredited by AAALAC International. For *M. tuberculosis* (*Mtb*) infection, animals were infected in an aerosol chamber with either standard “low-dose” inoculum of log phase Erdman strain *Mtb* at a concentration of 8*10^6^/mL in 5 mL of deionized water or “ultra low-dose” inoculum at a concentration of 6*10^5/mL in 5 mL of deionized water.

Animals were contained in aBSL3 biocontainment racks were given food and water ad libitum. All housing of and procedures involving animals were done according to the Animal Welfare Act, the Guide for the Care and Use of Laboratory Animals, the AVMA Guidelines on Euthanasia, and other federal statutes. All procedures were approved by the MSKCC Animal Care and Use Committee under approved animal study proposal 01-11-030.

### MAIT cell priming by intranasal inoculation

Mice were intranasally inoculated with 16.7 ug Pam2Cys (Invivogen) or Pam2Cys + 2 mM 5-A-RU (Aubé laboratory) + 50 uM methylglyoxal (Sigma) in 200 uL on Day 0, followed by 1 X PBS or 2 mM 5-ARU/50 uM methylglyoxal on days 1, 2 and 4. 5-ARU was synthesized as previously described (26), resuspended in sterile water, and cryopreserved in 12.5 mM aliquots. 5-A-RU was thawed as needed and directly added to methylglyoxal at the time of intranasal inoculation.

### Processing of tissues, in vitro stimulations and flow cytometry

Tissues were resected and underwent three minutes of bead beating at speed 6 using 10 2.0 mm zirconium oxide beads/sample in 1 mL (lungs and spleen) or 5 beads/sample in 0.5 mL of RPMI/10% fetal bovine serum in a Bullet Blender (Next Advance). Homogenized tissues were passed through 100 um strainers, centrifuged for 400XG for 3 minutes, supernatant was decanted and were then incubated for 5 minutes in RBC lysis buffer (Gibco ACK Lysing buffer) prior to final washing and in vitro assays or flow cytometry.

Lungs and spleen for *Mtb* culture were homogenized by four minutes of bead beating at speed 8 in 1 mL of 1x PBS/0.05% Tween 80. All CFU measurements were serial dilutions on 7H10 Middlebrook agar plates (Difco™) supplemented with OADC.

For in vitro re-stimulation assays, single cell suspensions of lung were incubated for 15 hours in media or 2 μM 5-A-RU added to 50 μM methylglyoxal in 96 well plates incubated at 37°C 5% CO_2_. Brefeldin A was added during the last two hours of stimulation.

Prior to fluorescent staining, FcγR blockade (CD16/CD32; clone 93) was performed for 15 minutes at room temperature in 50 μL. Extracellular staining was performed for 15 minutes at room temperature in 50 μL and intracellular staining was performed after 40 minutes of permeabilization/fixation at 4°C. For ESAT-6 and TB10.4 tetramers, cells were incubated with tetramers alone first for 1 hour at 37°C prior to staining with other antibodies. Intracellular antibody cocktails were incubated for 1 hr at 4 °C in 50 uL. We used MR1-5-OP-RU tetramers to identify MAIT cells and MR1-6FP tetramers as staining controls. ESAT-6 and TB10.4 tetramers were employed to identify *Mtb*-antigen specific T cells. All tetramers were produced by the NIAID Tetramer Core Facility (Emory University, Atlanta GA) (26). We used the following fluorochrome-labeled antibodies with their respective clones in parentheses: TCRβ(H57-597), FoxP3 (FJK-16S), MHC II (M5/114.15.2), CD25 (PC61), CD4 (GK1.5), CD69 (H1.2F3), IFNγ (XMG1.2), IL17A (TC11-18H10), F4/80 (BM8), CD45R/B220 (RA3-6B2), CD44 (IM7), granzyme B (QA16AA02), CD90.2 (30-H12), TCRγδ(GL3), CD19 (6D5), CD8 (53-6.7), PD-1 (29F.1A12), and NK1.1 PK136). All Mtb-infected tissues were fixed in 2% formaldehyde for two hours prior to acquiring on a BD Fortessa. Absolute numbers were normalized using Precision Count Beads™ (Biolegend). Flow cytometry data was analyzed using FCS express software (v. 7, De Novo Software, Pasadena, CA).

### Statistical analysis

All statistical analysis was performed in Prism (v. 8, GraphPad software, La Jolla, CA). Two-tailed unpaired t test with significance level of p<0.05 was used to compare immunologic or cfu differences between P2C/5-OP-RU or P2C treatment groups. A log-rank test with significance level of p<0.05 was used to compare survival in iNOS2 deficient mice.

## Supplementary information

**Supplemental Figure 1.**
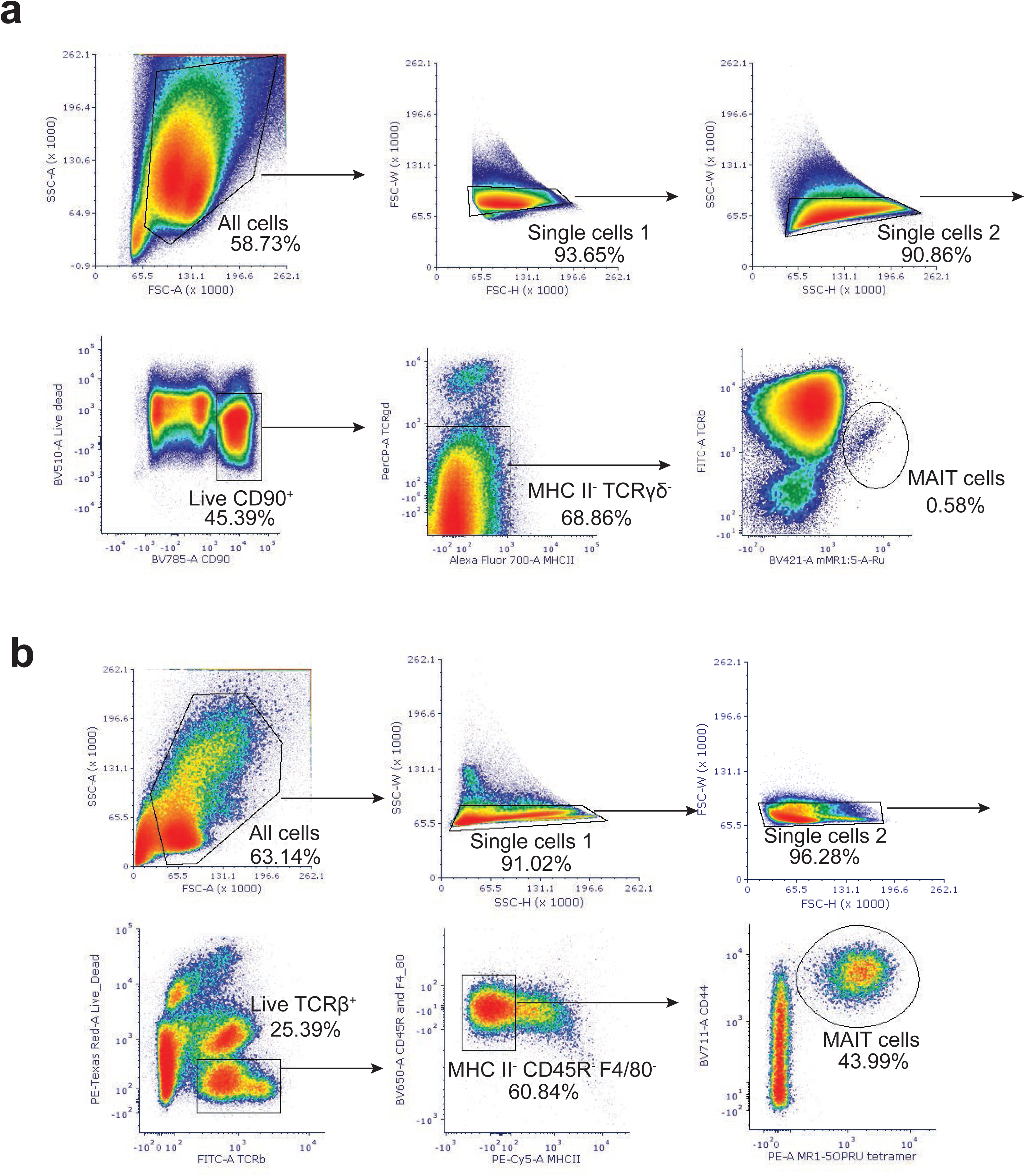
MAIT cell gating strategy in mice. **(a)** MAIT cell gating strategy for Figure 1. **(b)** MAIT cell gating strategy for Figures 2-5.

**Supplemental Figure 2.**
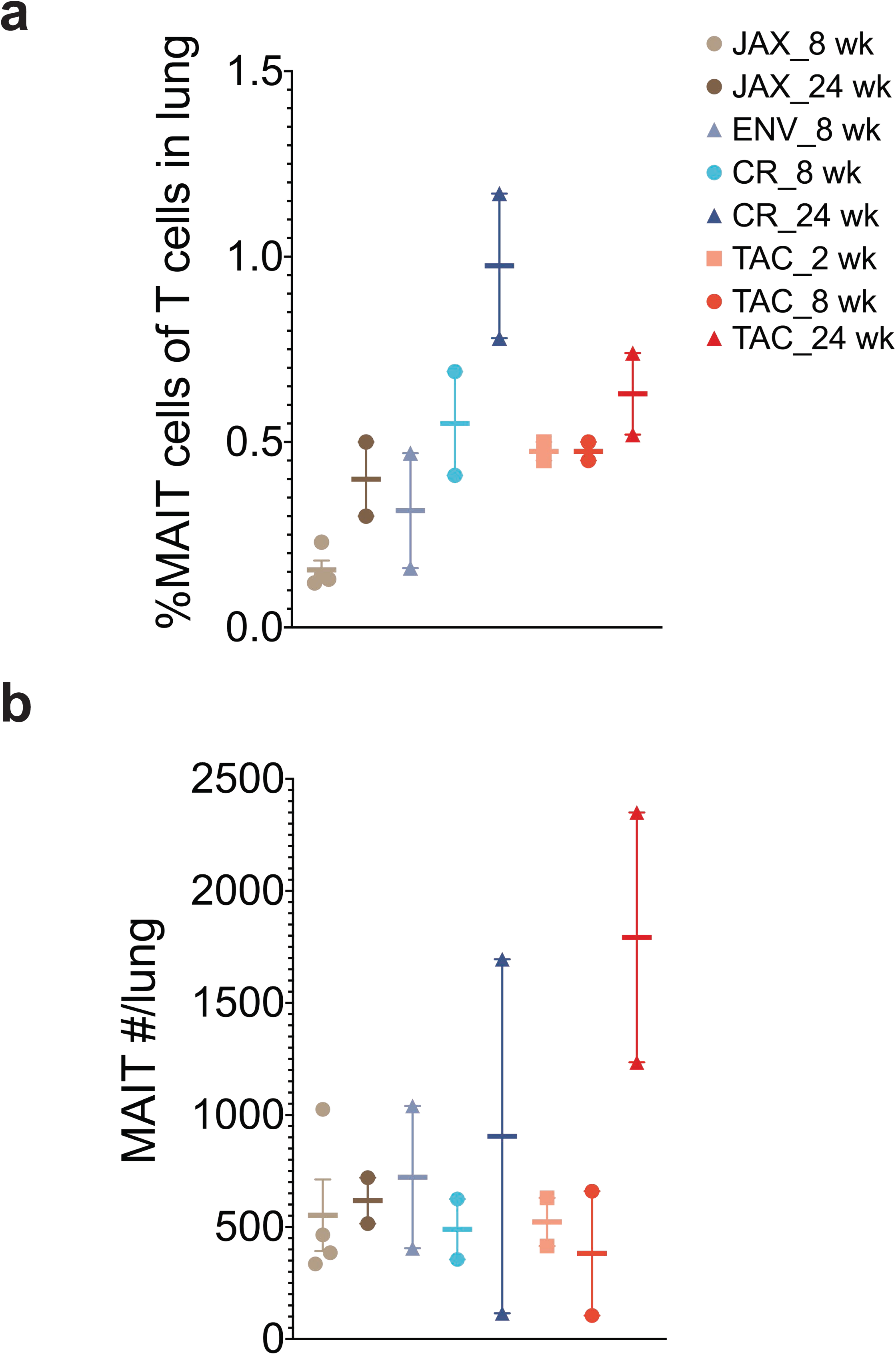
MAIT cells in mice from different breeders. **(a)** Mean% MAIT cells of CD90^+^ cells and **(b)** mean MAIT cell absolute number in mice from different vendors. n=2-4 mice per group. All mice were C57Bl/6 strains from the following vendors: JAX: Jackson; ENV: Envigo; CR: Charles River; TAC: Taconic; wk=weeks

**Supplemental Figure 3.**
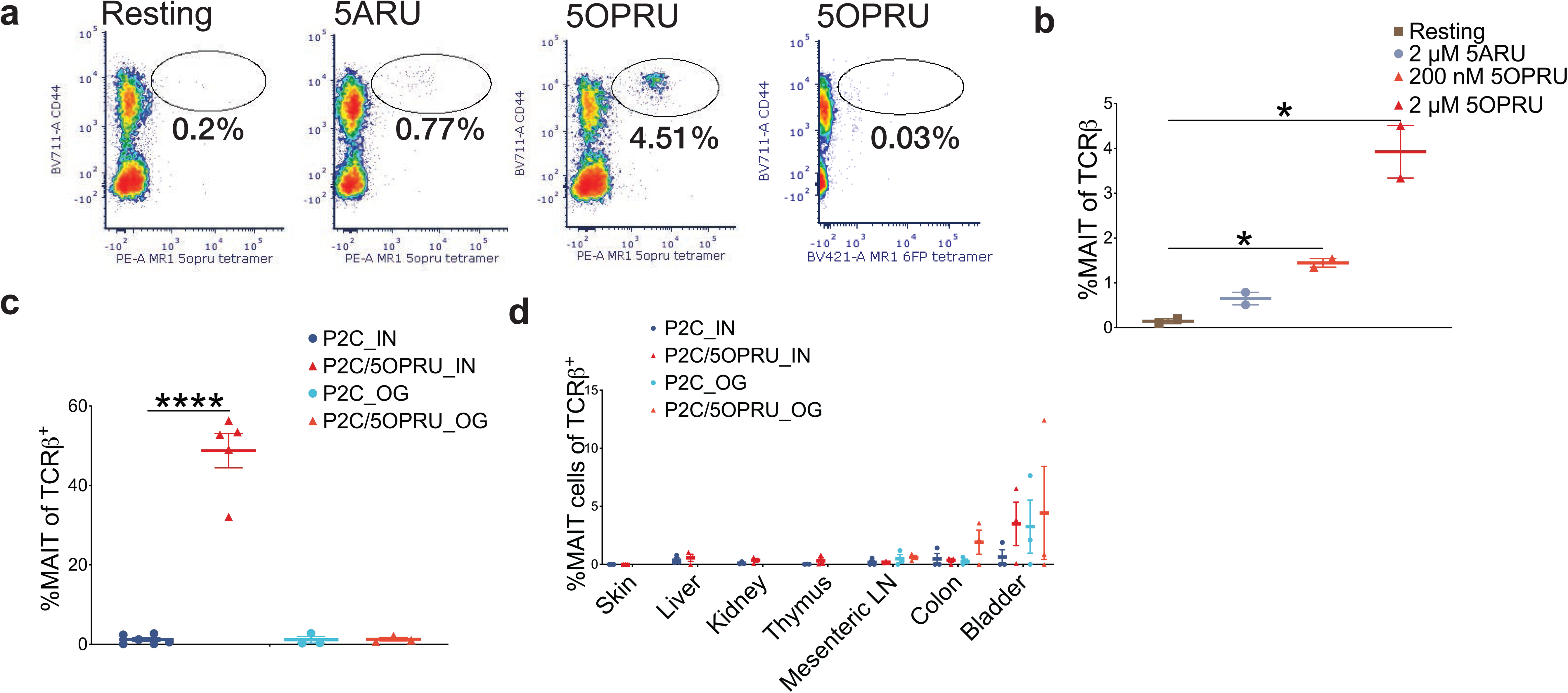
Murine MAIT cell expansion in vitro and in vivo. **(a)** Representative flow cytometry density plots from splenocytes from three mice after 7 days of in vitro incubation under different stimulation conditions. Plots demonstrate identification of splenic MAIT cells by MR1-5-OP-RU tetramers. Staining controlled with MR1-6FP tetramers. **(b)** Mean % MAIT cells of T cells +/− SEM after seven days of in vitro incubation. **(c)** Mean %MAIT cells of T cells in lungs of mice 7 days after priming by intranasal (IN) inoculation or oral gavage (OG). n=3-5 mice/group. **(d)** Mean% MAIT cells of T cells +/− SEM in various tissues 14 days after IN or OG priming. n=3-6 mice/group. Each panel is representative of at least two independent experiments. Statistical analyses were performed using unpaired t-tests. *p<0.05 **p<0.005

**Supplemental Figure 4.**
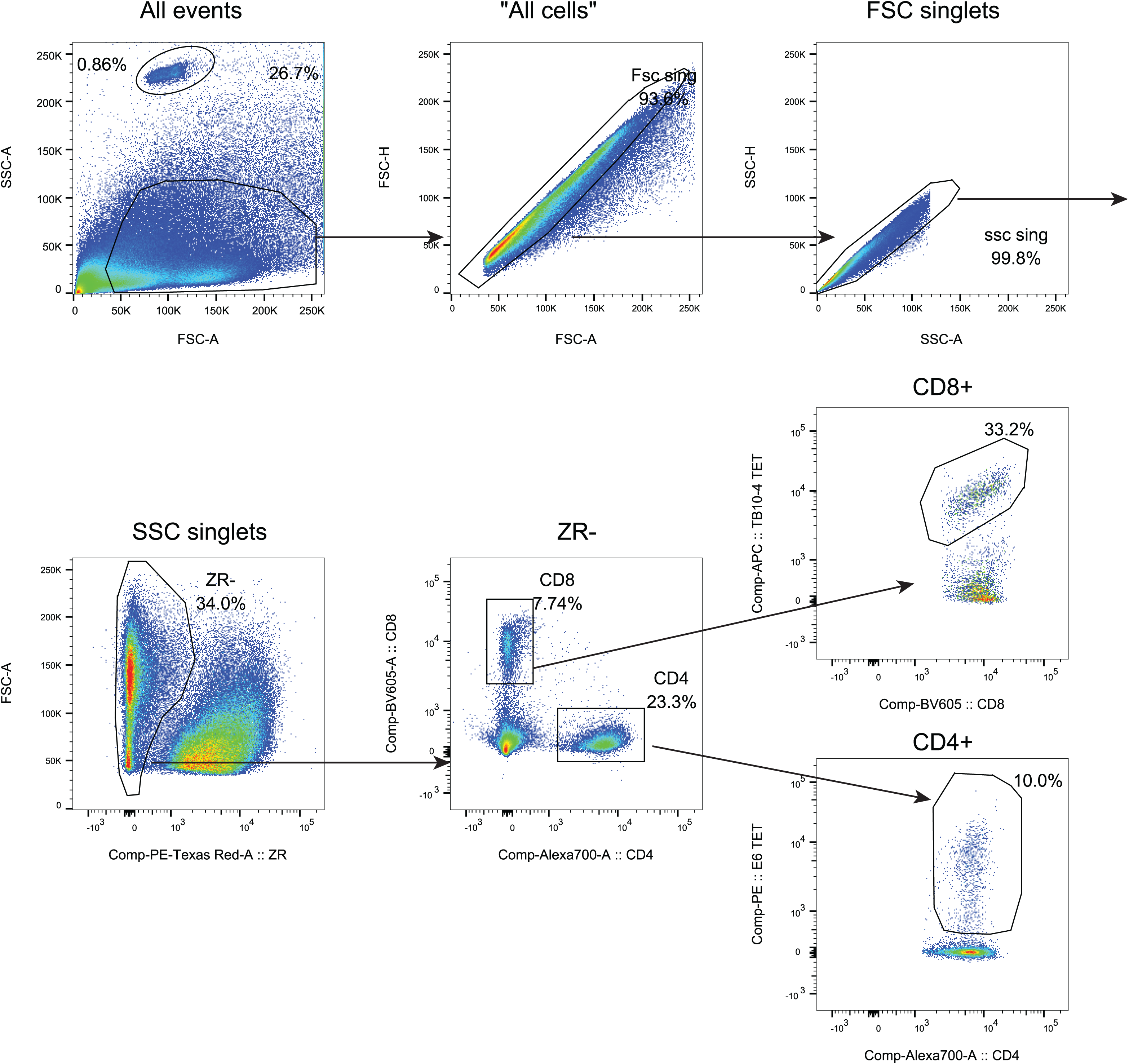
ESAT-6 and TB10.4 tetramer gating strategy. Gating strategy to identify ESAT-6- or TB10.4-specific T cells among CD4^+^ or CD8^+^ T cells, respectively using tetramers.

**Supplemental Figure 5.**
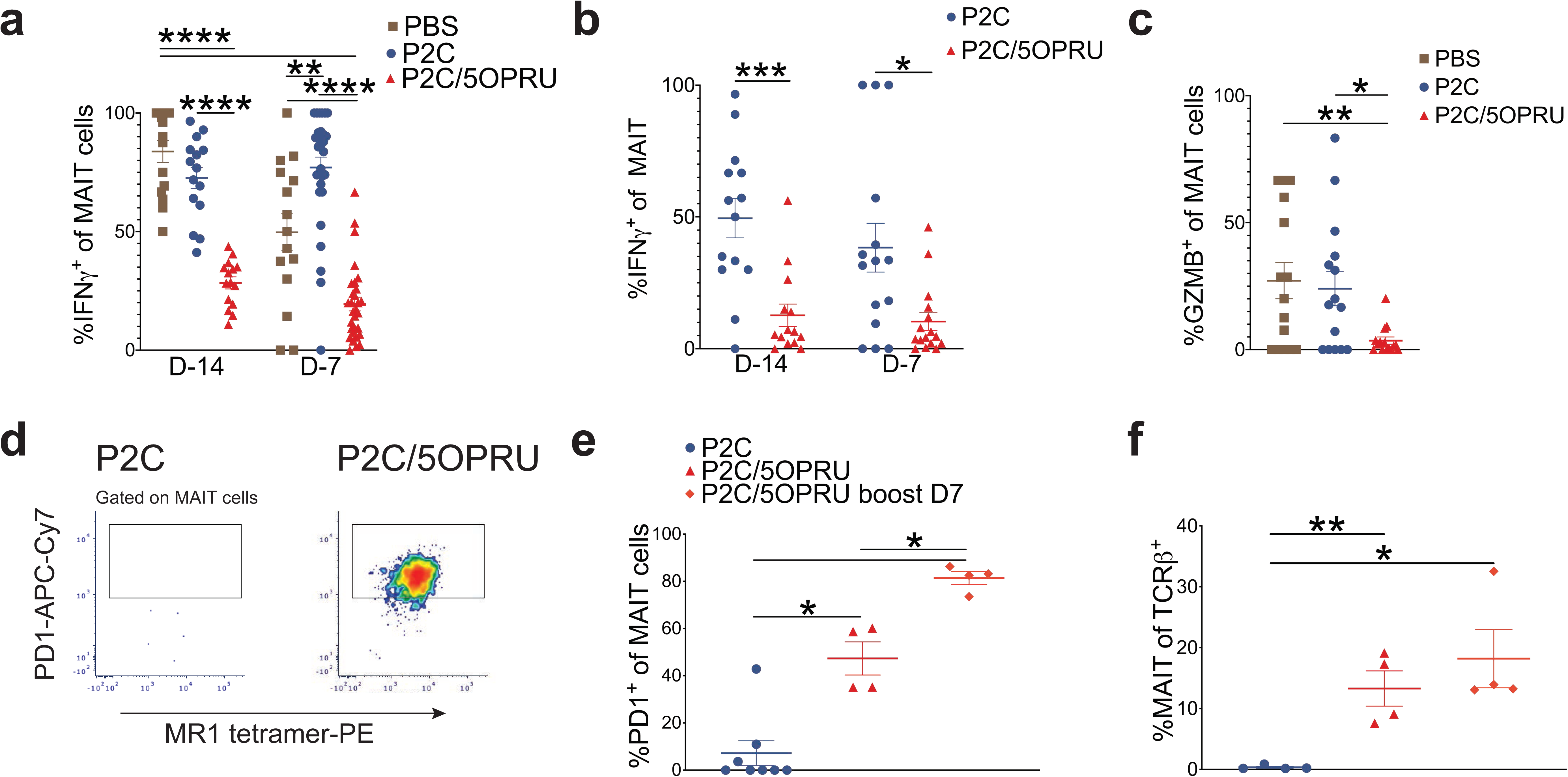
MAIT cells are driven towards exhaustion during chronic activation. **(a, b)** Mean % IFNγ^+^ MAIT cells +/− SEM at day 14 (a) and 24 (b) post-infection stratified by infection schedule **(c)** Mean% granzyme B^+^ (GZMB) MAIT cells +/− SEM at 14 days post-infection. n= **(d)** Representative flow cytometry density plots in two mice demonstrating PD1 staining in MAIT cells 14 days after priming. **(e)** Mean % PD1^+^ of MAIT cells +/− SEM after 14 days of priming compared to mice that received additional priming with P2C/5-OP-RU on Day 7 (boost). **(f)** Mean % MAIT cells of T cells +/− SEM in the same conditions as panel e. n=15-30 mice/group. Data in panels a and b represent 2 independent experiments. Data in panel c-f represent one independent experiment. Statistical analyses were performed using unpaired t-tests. *p<0.05 **p<0.005 *** p<0.0005 ****p<0.0001

